# Activation of the mTOR/ Akt pathway in thymic epithelial cells derived from thymomas

**DOI:** 10.1101/317297

**Authors:** Jean-Michel Maury, Claire Merveilleux du Vignaux, Gabrielle Drevet, Virginie Zarza, Lara Chalabreysse, François Tronc, Nicolas Girard, Caroline Leroux

## Abstract

The pathogenesis of thymic epithelial tumors remains poorly elucidated. The PI3K/Akt/mTOR pathway plays a key role in various cancers; interestingly, several phase I/II study reported a positive effect of mTOR inhibitors in disease control in thymoma patients. A major limit for deciphering cellular and molecular events leading to the transformation of thymic epithelial cells or for testing drug candidates is the lack of reliable in vitro cell system

We analyzed protein expression and activation of key players of the Akt/mTOR pathway namely Akt, mTOR, and P70S6K in thirteen A, B and AB thymomas as well as in normal thymuses. While only Akt and the phospho-Akt were expressed in normal thymuses, both Akt and mTOR were activated, with B2 thymomas expressing higher level of activated phospho-Akt than A or AB subtypes. Phospho-P70S6K was expressed in all thymic tumors whatever their subtypes, and absent in normal thymus. Interestingly, in primary thymic epithelial cells maintained for short period of time after their derivation from seven AB and B thymomas, we report the activation of Akt; mTOR and P70S6. Finally, we analyzed the effect of mTOR inhibitor on thymoma derived epithelial cells and showed that rapamycin (100 nM/ ml) significantly reduced cell proliferation.

Our results suggest that the activation of the Akt/ mTOR pathway might participate to the cell proliferation associated with tumor growth. Ultimately, our data enhance the potential role of thymic epithelial cells derived from tissue specimens for *in vitro* exploration of molecular abnormalities specific to rare thymic tumors.

## Introduction

Thymic epithelial tumors (TETs) are rare epithelial malignancies (0.2-1.5%) of the anterior mediastinum, with an estimated incidence about 1.3-3.2 cases worldwide [1]. The WHO classification distinguishes thymomas and thymic carcinomas [2]. Thymomas are defined as A, AB, B1, B2, B3 sub-types according to the morphology of tumor epithelial cells, the proportion of non-tumoral thymic lymphocytes (decreasing from B1 to B3) that are associated with tumor cells, and their similarities to normal thymic architecture. Thymic carcinomas present with a high degree of epithelial cells atypia associated with a loss of normal thymic architecture.

Surgical resection is the corner stone of the multimodal treatment of thymomas [3]. Tumor stage [4] and radical complete surgical resection have been shown as independent prognosis factor of best outcome [5-7]. Advanced or metastatic cases are treated with induction chemotherapy, surgery, combined radiation-chemotherapy [8-11] with variable outcomes from 12 to 60% response [12-15].

The pathogenesis of thymic epithelial tumors remains poorly elucidated. Sustained efforts have been made to characterize molecular abnormalities occurring in TETs to improve their treatment and eventually the patient prognosis. Sequencing of 197 cancer-related genes revealed the presence of non-synonymous somatic mutations in over 60% thymic carcinomas and barely 15% thymomas [16]. The most frequent mutations (26 % of thymic carcinomas) were located in the p53 tumor suppressor gene [17]. The Cancer Genome Atlas recently reported results using multi-platform omics analyses on 117 TETs, leading to define four subtypes defined by genomic hallmarks [17]; GTF2I was confirmed as an oncogene associated with type A thymoma, and mutations in HRAS, NRAS, and TP53 were identified in thymomas. A major limit of those studies was the use of tumor tissue specimens precluding specific analysis of epithelial tumor cells while lymphocytes may present with a high level of expression of genes related to carcinogenesis [17-23]

The PI3K/Akt/mTOR pathway plays a key role in various cancers and among them thymic tumors. Mutations of genes encoding regulatory subunit of PI3K were recently reported in a tumorigenic thymic carcinoma cell line, using targeted exome sequencing, predicting the efficacy of PI3K inhibitors [24]. Several phase I/II study of mTOR inhibitors were reported in advanced thymic epithelial tumors, reporting on high disease control rates [25-27]. Meanwhile the cellular dysregulation of the Akt/ mTOR pathway has not been described in thymomas. Using primary thymic epithelial cells derived from AB and B thymomas, we report the dysregulation of the Akt/ mTOR pathway in thymomas and the anti-proliferative effect of rapamycin on thymic epithelial cells.

## Materials and Methods

### Biological samples

Between January 2015 and December 2017, thymic samples from patients that have undergone removal surgery for thymic tumors (N=13) (Table 1) or cardiac surgery (N=2) for normal thymuses have been included in our study. Tumoral and normal tissues and their associated data (Table 1) were obtained from the CardioBiotec biobank (CRB-HCL, Hospices Civils de Lyon BB-0033-0046), a center for biological resources authorized by the French ministry of social affairs and health. Patients with thymic epithelial tumors were identified by the department of pulmonary medicine and thoracic oncology (Groupement Hospitalier Est, HCL Lyon). All samples used in the study were collected and used in accordance with the ethical rules of the bio bank and in agreement with the French legislation. All patients signed a written informed consent.

**Table 1.** Clinical features of thymomas included in the study. ^1^ Subtype has been defined according to the WHO classification, ^2^ staging according to the Masaoka-Koga classification, ^3^ TNM according to the ITMIG guidelines. ^4^ unknown, pleural metastasis; ^5^ received Caelyx therapy for Kaposi treatment, ^6^ received cisplatin, adriamycin and cyclophosphamide neo-adjuvant treatment.

| Patient # | Age (yr) | Sex | Myasthenia gravis | Tumor size (cm) | Previous chemotherapy | Subtype<sup>1</sup> | Staging<sup>2</sup> | TNM<sup>3</sup> |
| --- | --- | --- | --- | --- | --- | --- | --- | --- |
| 3153 | 73 | M | No | ?<sup>4</sup> | No | A (pleural metastasis) | IVA | T3N0M1a |
| 3154 | 70 | M | No | 5.7 | No | A (Micronodular with lymphoid stroma) | IIA | T1aN0M0 |
| 2635 | 57 | M | No | 6 | No | AB | I | T1aN0M0 |
| 2646 | 83 | F | No | 5 | No | AB | I | T1aN0M0 |
| 2836 | 46 | M | No | 10 | No | AB | I | T1aN0M0 |
| 3147 | 70 | M | No | 11 | No | AB | III | T2N0M0 |
| 3148 | 65 | F | No | 3.5 | Yes<sup>5</sup> | AB | I | T1aN0M0 |
| 3152 | 68 | F | No | 3.5 | No | AB | I | T1aN0M0 |
| 2779 | 38 | M | No | 6.5 | No | B2 | IIA | T1aN0M0 |
| 3146 | 71 | M | Yes | 3.2 | No | B2 | IIA | T1aN0M0 |
| 3149 | 63 | F | No | 9 | Yes<sup>6</sup> | B2/ B3 | I | T1aN0M0 |
| 3150 | 66 | M | Yes | 6 | No | B2/ B3 | III | T3N0M0 |
| 2637 | 68 | M | No | 7 | No | Thymic carcinoma | III | T3N0M0 |

Immediately after surgery, the thymic tumors were placed in RPMI 1640 cell medium supplemented with penicillin and streptomycin and processed in a biosafety level 2 lab in the next two hours for cell derivation or frozen in liquid nitrogen for further use.

### Derivation of thymic epithelial cells

Thymic epithelia cells (TECs) were obtained from thymic epithelial tumors as previously described with minor modifications [28, 29]. Thymic tissues were immediately placed in ice cold RPMI 1640 medium, cut in 1-3 mm^3^ pieces and transferred in “Liberase^™^ (Roche) digestion solution (RPMI 1640 medium supplemented with 0.5 U/ml of Liberase^™^ and 0.1% w/v DNase I) with ~ 2 ml of digestion solution/ cm^3^ of tissues. After 20 min incubation at 37°C under gentle agitation, supernatants were collected, mixed (v/v) with 1X PBS supplemented with 0.1 % bovine serum albumin, and 0.5 mM EDTA, and centrifuged at 480g for10 min at 4°C. The digestion procedure was repeated 4-6 times. Cells were counted in trypan blue using a Cellometer (Nexcelon BioSciences), seeded at 2-4.106 cells/ cm2 in “TEC medium” (RPMI 1640 medium supplemented with 2% Ultroser^®^ serum substitute (Pall corporation) and penicillin/streptomycin) and incubated at 37°C, 5% CO_2_ in humid atmosphere. After 24 hours, cell culture supernatants were centrifuged at 480g for10 min at 4°C to eliminate non-adherent cells, supplemented with an equal volume of new “TEC medium” and add to the cultured cells. Cells were checked daily and passaged at confluence with trypsin EDTA solution.

### Phenotypic characterization of primary thymic epithelial cells

Primary thymic epithelial cells were cultured on treated glass slides (LabTekTM, Thermo Scientific) in “TEC medium”, rinsed in PBS and fixed with ice cold acetone. After 20 min rehydration in PBS, cells were incubated for one hour at room temperature with BM4048 mouse monoclonal anti-cytokeratin (Acris) or mouse monoclonal anti-vimentin (Sigma) diluted in PBS supplemented with 1% bovine serum albumin as recommended. After PBS washes, cells were incubated for one hour at room temperature with anti-mouse IgG Dylight antibodies Eurobio). Nuclei were stained with DAPI for 10 min and mounted with Fluoromount G (Electron Microscopy Sciences). Microscopic examinations were performed using an Axio-Imager Z1 epifluorescence microscope and analyzed using the Zen software (Zeiss).

### Protein expression in thymic tissues and thymic epithelial cells

Proteins were prepared from thymic tissues and primary thymic epithelial cells using respectively T-PER^®^ extraction reagent or M-PER^®^ extraction reagent (Thermo Fisher) supplemented with “HALT protease and phosphatase inhibitor cocktail^®^” (Thermo Fisher) Lysates were homogenized by sonication on ice and proteins were measured from the collected supernatants. Twenty to thirty micrograms of total proteins were separated on SDS-PAGE, transferred onto PVDF membrane and used for the detection of Akt with “Akt1 Precision Ab^™^ antibody” (BIORAD VMA00253), phospho-Akt using “phospho-Akt (Ser 476) antibody” (Cell signaling, 9271S), mTOR using “mTOR PrecisionAbTM Antibody” (BIORAD, VPA00174), phospho mTOR using “phospho-mTOR (Ser2448) Antibody”; (Cell signaling, 2971), phospho p70S6k using “phospho-p70 S6 Kinase (Thr389) Antibody” (Cell signaling, 9205) and β-actin (Monoclonal Anti-β-Actin–Peroxidase antibody, Sigma) as recommended. Immunoreactive bands were detected with goat anti-rabbit IgG (whole molecule) - peroxidase antibodies produced in goat (Sigma) and revealed using the “Clarity Max^™^ Western ECL Blotting Substrate” (BIORAD) on a “ChemiDoc^™^ Imaging System” (BIORAD).

### Thymic epithelial cells proliferation upon rapamycin exposure

Primary thymic epithelial cells (5000 cells/well) derived from thymomas #3146, #3147 and #3149 were incubated in “TEC medium” in 96 well plates for twelve hours then treated with 1, 10 or 100 nM rapamycin (Sigma). Controls with DMSO or untreated cells have been included. Proliferation was measured 24, 48 and 72 hours after rapamycin treatment with the “CellTiter-Glo^®^ Luminescent Cell viability assay” (Promega). All tests have been repeated at least twice and performed in triplicates.

### Sequencing mutation of the PI3K complex: PI3KCA and PI3KR1and GTF2I

Mutations have been sought using primers designed in exons 1 to 21 of the PI3KCA gene (GenBank NM06218.3), exons 5 to 16 of the PI3KR1 (GenBank NM181523) gene and have been sought in 5-10 and exons 10 to 15 of the GTF2I (GenBank NM001518.4) gene (Table 2).

**Table 2.**
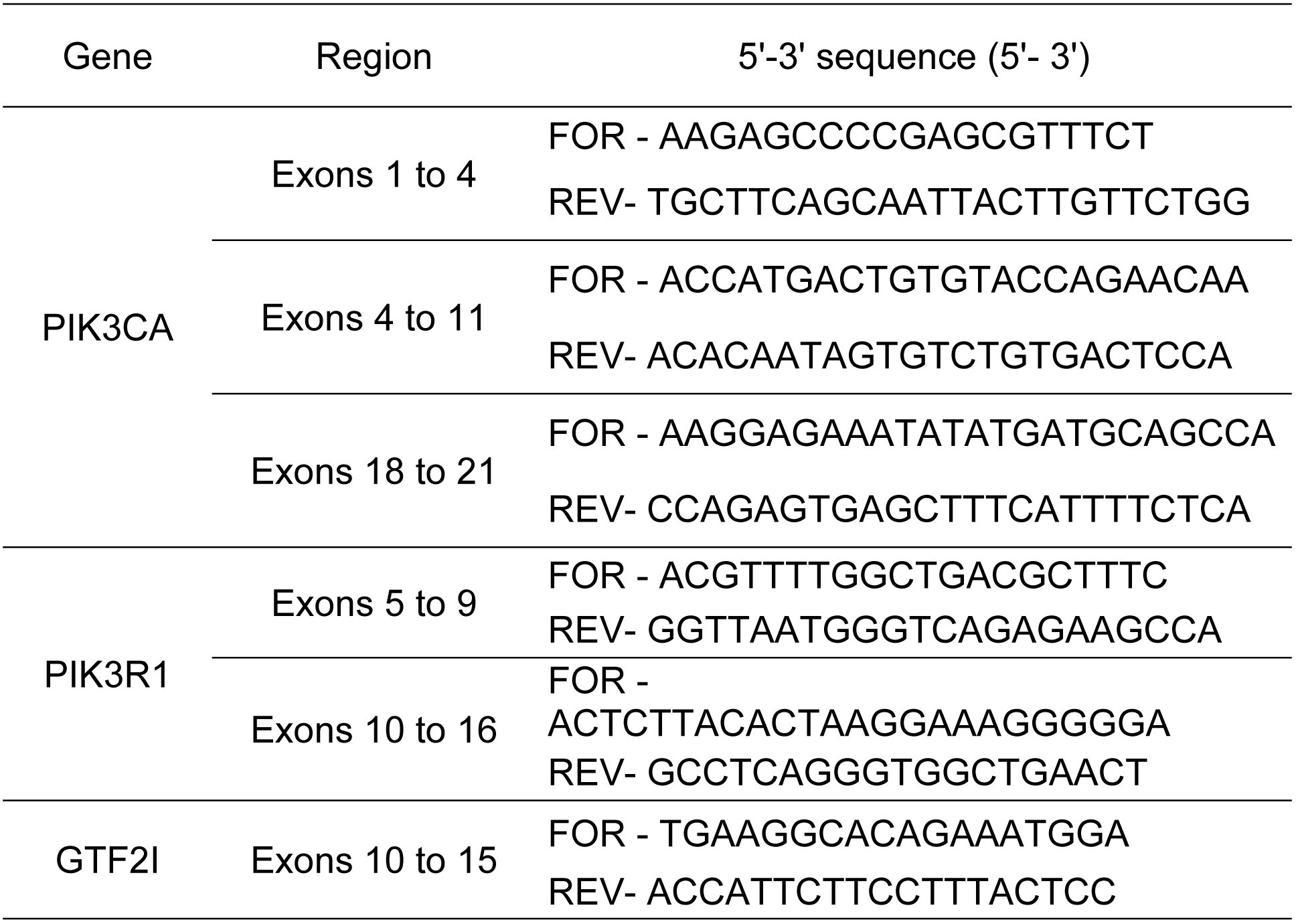
Primers used to amplify GTF2i and PIK3 genes.

Primers were selected with PrimerBlast software. Briefly, RNAs were extracted with the “Pure Link RNA minikit” (Ambion), reverse transcribed using “iScript cDNA synthesis kit” (Bio-Rad) and amplified with “KAPA HIFI hotstart polymerase” (Clinisciences) for 35 cycles (2 min at 98°C, 15 sec at 60°C and 30 sec at 72°C) on a Mic qPCR system (BioMolecularSystem). Amplicons were controlled on agarose gel, sequenced (GTAC Biotech) and analyzed with the Vector NTI software (Invitrogen). The potential impact of discovered mutation was analyzed with the “PolyPhen-2 (Polymorphism Phenotyping v2) predictive model”.

### Statistical analysis

Statistical analyses (threshold of α = 0.05) were performed using the t test with the “GraphPad Prism” (GraphPad) software. All tests were done with a significant threshold of α = 0.05.

## Results

### Clinical features

From January 2015 to December 2017, thirteen patients (9 males and 4 females, mean age of 64.46 ± 11.72 years (38 – 83)) who have undergone surgery (Table 1) for thymic tumors (3154, 2635, 2646, 2836, 3147, 3148, 3152, 2779, 3146, 3149, 3150, 2637) or for pleural relapse (3153) of a type A thymoma removed 25 months before have been included in our study (Table 1).

Among them, patients 3148 and 3149 received chemotherapy before surgery respectively to treat cutaneous Kaposi lesions with Caelyx and initially locally advanced thymoma with cisplatin, adriamycin and cyclophosphamide. Two patients (3146 and 3150) presented myasthenia gravis and were treated with intra venous polyvalent immunoglobulins one week before surgery. According to the WHO pathological classification, tumors were of type AB for 6 patients (2635, 2646, 2836, 3147, 3148, and 3152), B2 for 4 patients 2779, 3146, 3149, 3150 and thymic carcinoma for patient 2637. Subtype A thymic samples were from pleural metastasis in patient 3153 and micronodular thymoma with lymphoid stroma, a rare presentation of type A thymoma for patient 3154. According to the Masaoka classification, tumors were of stage I for six patients, IIA for two patients, III for three patients and IVA for one patient.

### Derivation of thymic epithelial cells

Primary thymic epithelial cells were successfully derived from the thirteen samples immediately after tumor removal and expanded in culture. Daily observation under phase microscope showed that cells had an epithelioid morphology (Fig 1). Cells proliferated, and the epithelial morphology was maintained along the study up to passages 6 to 7, corresponding to a median time in culture of 56 days.The primary thymic cells derived from the tumors expressed cytokeratin (Fig 1), a marker of epithelial cells, with a mean of 79% positive cells except for cells derived from the thymic carcinoma 2637 that were negative for cytokeratin but expressed vimentin.

**Fig 1.**
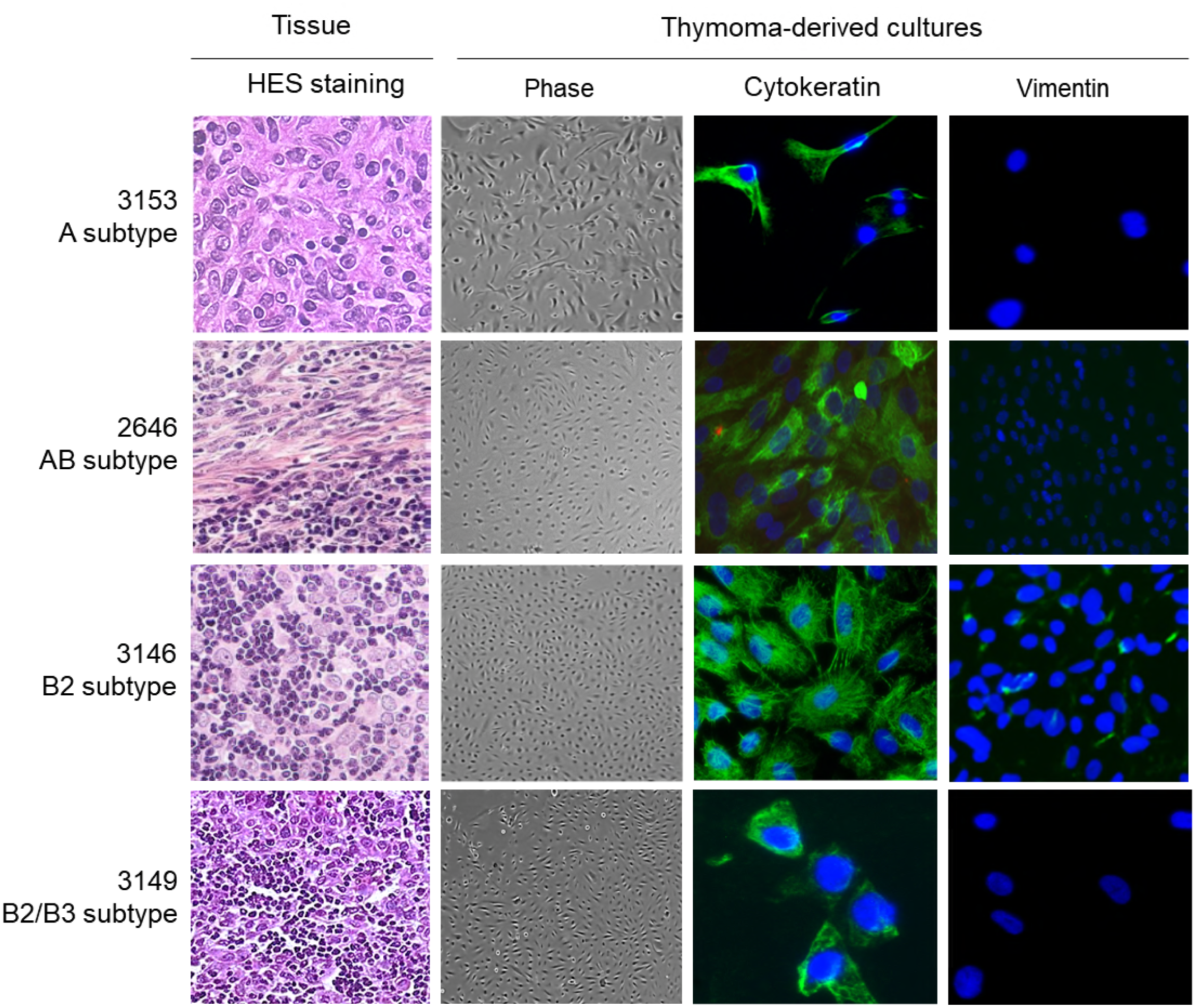
Primary thymic epithelial cells derived from A, AB and B thymomas. Representative **c**ell cultures derived from patients 3153 (pleural metastasis of A thymoma), 2646 (AB thymoma), 3146 (B2 thymoma) and 3149 (B2/ B3 thymoma). HES (hematoxylin-eosin-saffron) staining were used to characterize the thymic tissues. Thymoma-derived cells were observed daily by phase microscopy and stained for their expression of cytokeratin and vimentin. Nuclei stained with DAPI (blue).

To summarize, we successfully derived primary epithelial cells from thymic tumors and pleural metastasis of a type A thymoma. These cells expressed cytokeratin and were able to proliferate *in vitro* over several passages. The subsequent analyses (detection of protein as well as proliferation studies) were performed on early passages, to be as close as possible to the *in vivo* phenotype.

### Mutations of Pi3K and GTF2i genes

A screening of PI3K and GTF2I mutations was performed on all tumors. Among the 12 tumors, only patient 3149 (B2/ B3 thymoma) carried a non-conservative A/C transversion localized on position 56 of exon 2 of PI3KCA (Fig 2A). The mutation induced a K → Q amino-acid change, predicted as deleterious with the PolyPhen-2 prediction. We also detected a GTF2I mutation for tumor 3154, reported as a micronodular thymoma with lymphoid stroma, a rare presentation of type A thymoma (Fig 2B). The mutation located on exon 15 was associated with a non-conservative T/A transversion leading to L → H amino-acid change in the deduced amino-acid sequence (Fig 2B).

**Fig 2.**
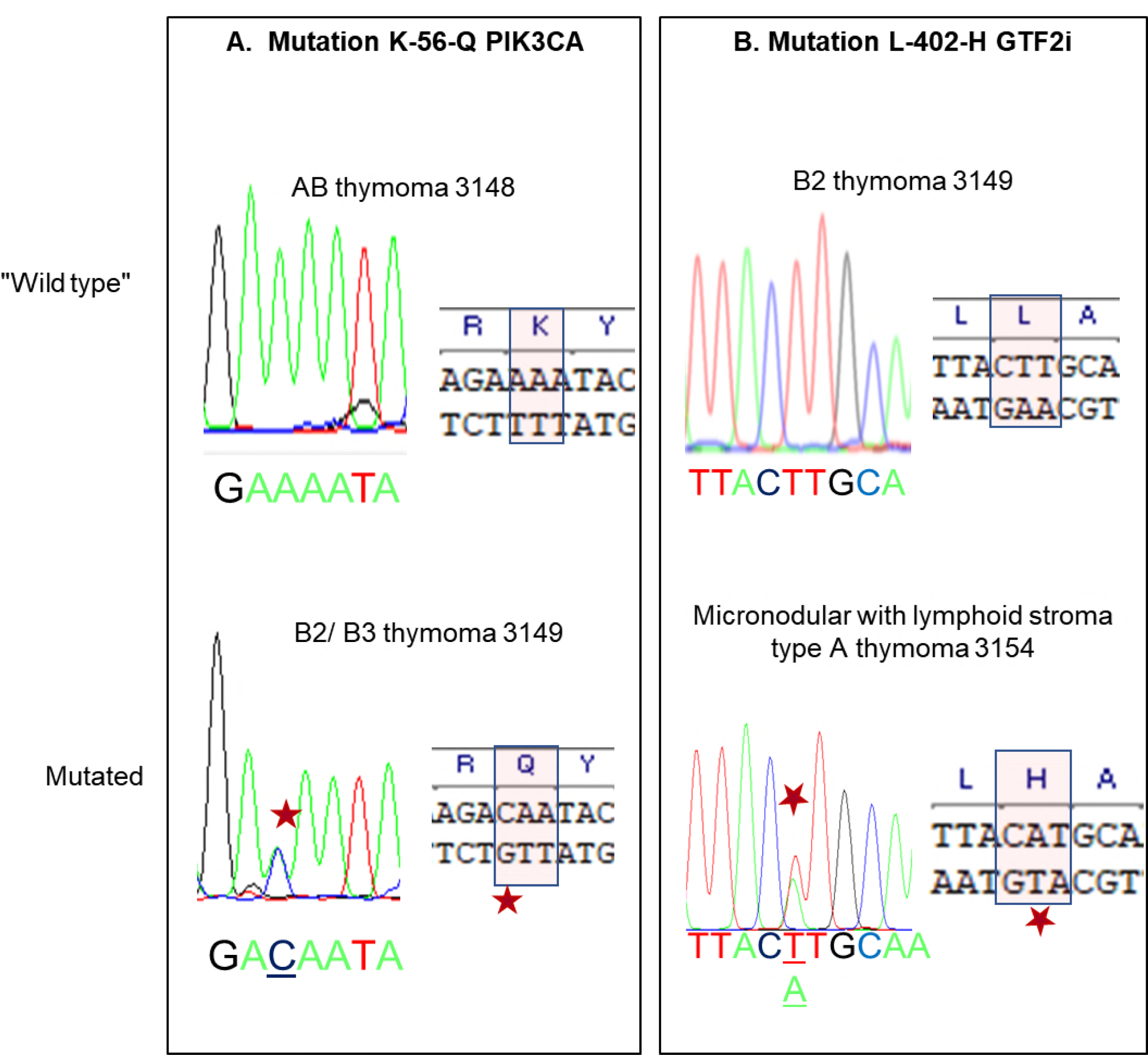
Mutations in PIK3CA and GTF2i. PIK3CA and GTF2i have been amplified from total RNA extracted from thymoma, sequenced and compared to reference sequences. A. C/A mutation in exon 2 of PIK3CA in B2/ B3 thymoma #3149. B. T/A mutation in exon 15 of GTF2i in micronodular with lymphoid stroma type A thymoma #3154

#### *A*ctivation of the Akt/ mTOR/ P70S6K pathway in thymomas

The Akt/ mTOR pathway is a key pathway implicated in cell proliferation. We analyzed its activation in thymomas as well as in normal thymuses from cardiac surgery. In these latter ones, only Akt and the activated phospho-Akt were expressed while mTOR, phospho mTOR or phospho P70S6K proteins were undetectable (Fig 3A). In thymomas, both Akt and mTOR were activated (Fig 3A), as shown by the detection of phosphorylated proteins. The protein signals have been quantified using ImageLab (Biorad) and expressed as the ratio [protein of interested/ β-actin] (Fig 3B). It suggested that B2 thymomas expressed higher level of Akt and activated phospho-Akt than A or AB subtypes (Fig 3B). A Mann and Whitney analysis showed a statistically significant overexpression of Akt in B2 *vs.* AB and A thymomas (respectively p=0.0017 and p=0.0095). Finally, phospho-P70S6K was expressed in all thymic tumors whatever their subtypes, and absent in normal thymus.

**Fig 3.**
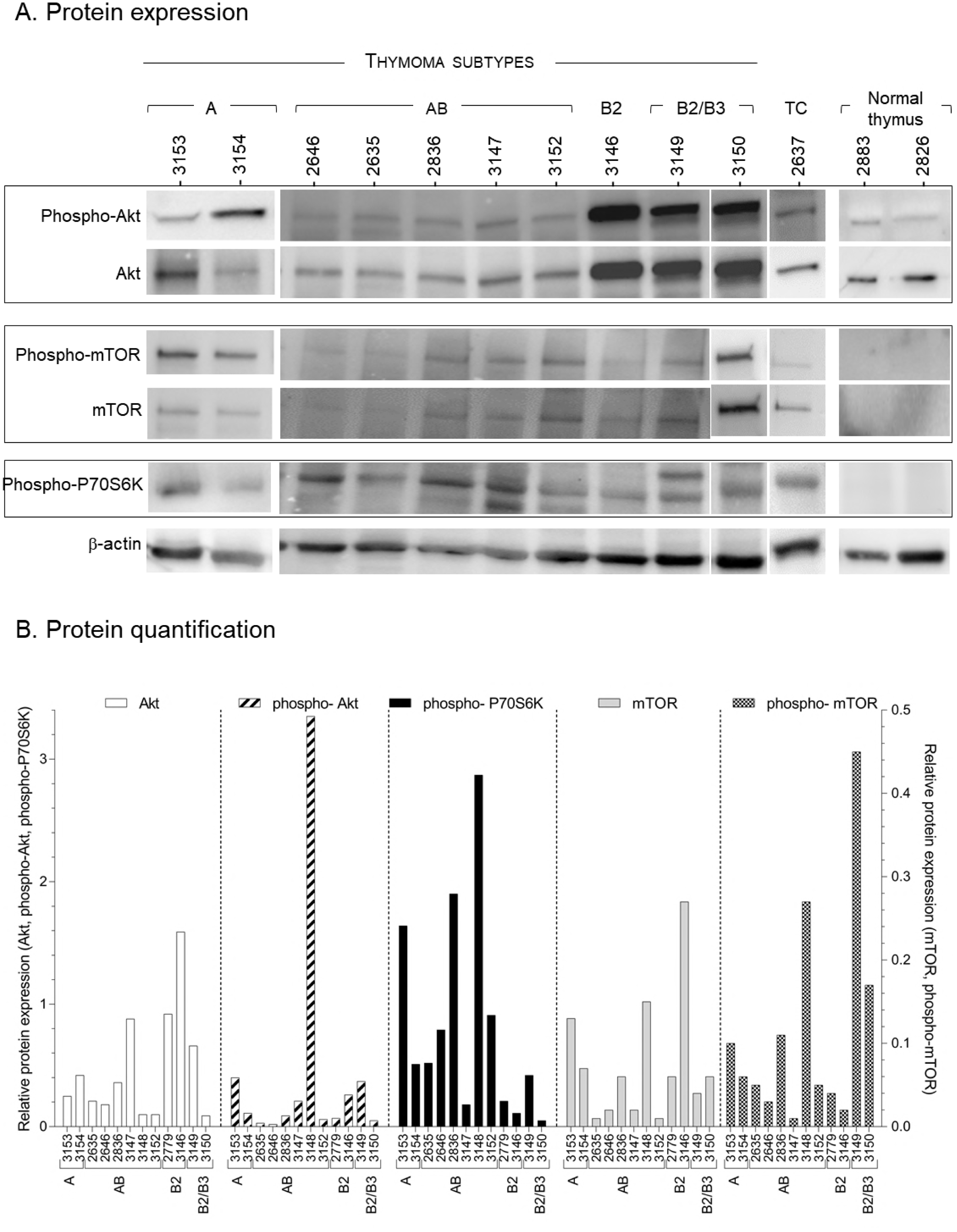
Activation of the Akt/ mTOR pathway in thymic epithelial tumors. A. Total and phosphorylated protein expression was analyzed using antibodies directed against total Akt and phosphorylated-Akt (phospho-Akt) (60 kDa), total mTOR and phosphorylated-mTOR (phospho-mTOR) (289 kDa), phosphorylated P70S6K (phospho-P70S6K) (70 kDa) and β-Actin (40 kDa). B. Protein expression has been measured and expressed as [protein of interest/ actin] relative expression in A, AB and B (B2 and B2/ B3) thymomas, thymic carcinoma (TC) or normal thymus. Detection have been repeated at least 3 times. Left Y axis: Akt, phospho-Akt, phospho-P70S6K expression; right Y axis: mTOR, phospho-mTOR.

Overall, the Akt/ mTOR pathway was activated in A, AB and B thymomas as demonstrated by the detection of phosphorylated Akt, mTOR and P70S6K proteins, with various level of expression that probably reflected the relative frequency of tumoral and non tumoral cells within the tumors.

### Activation of the Akt/ mTOR pathway in thymic epithelial cells derived from thymomas

To analyze the role of the Akt/ mTOR pathway in proliferation of tumoral thymic epithelial cells, we analyzed expression of mTOR, Akt and P70S6K in 7 thymoma-derived epithelial cells. Interestingly, all the primary cells expressed detectable levels of phosphorylated Akt and mTOR while phospho-P70S6K was barely detectable except in 3146-B2 thymoma (Fig 4). This suggests that the activation of the Akt/ mTOR pathway might participate to the cell proliferation associated with tumor growth.

**Fig 4.**
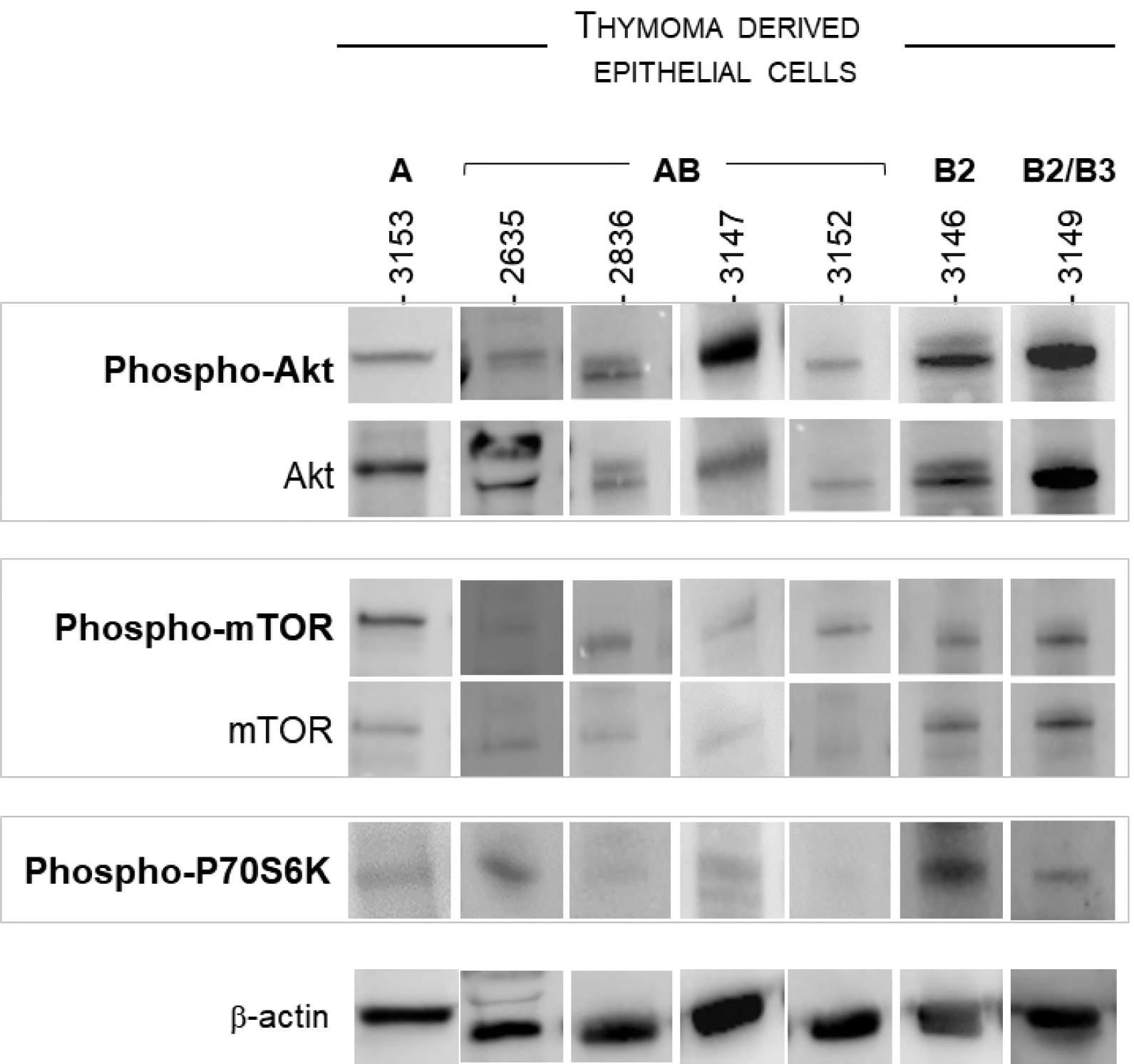
Activation of the Akt/ mTOR pathway in thymoma-derived thymic epithelial cells. Total and phosphorylated protein expression was analyzed using antibodies directed against total Akt and phosphorylated-Akt (phospho-Akt), total mTOR and phosphorylated-mTOR (phospho-mTOR), phosphorylated P70S6K (phospho-P70S6K) and β-Actin in thymic epithelia cells derived from A, AB or B2 thymomas.

### Effects of rapamycin treatment on tumoral TECs proliferation

We analyzed the effect of rapamycin, an inhibitor of mTOR, on cell proliferation of thymic epithelial cells. We focused our analysis on three cell cultures, at early passages, derived from AB (# 3147), B2 (# 3146) and B2/ B3 (#3149) thymomas because they had good proliferative abilities with ~24 hours doubling time (not shown) that is a prerequisite for studying rapamycin inhibition over a 72-hour period. Importantly, these three thymomas and their derived cells expressed mTOR and phospho-mTOR (Figures 3 and 4). The proliferation rate was measured daily, in triplicates and repeated at least twice over 72-hour treatment with 1 nM, 10 nM or 100 nM rapamycin. No effect was detectable with 1 nM rapamycin but with 10 or 100 nM, the proliferation was reduced in all three primary cell cultures, highly significantly with 100 nM (Fig 5) and cell death was detectable in the treated or untreated assays (not shown). In cell treated with 100 nM rapamycin, the proliferation rate was ~30% inferior to that of untreated cells; this reduction was similar to that observed in A549 cells treated with rapamycin used as control (not shown).

**Fig 5.**
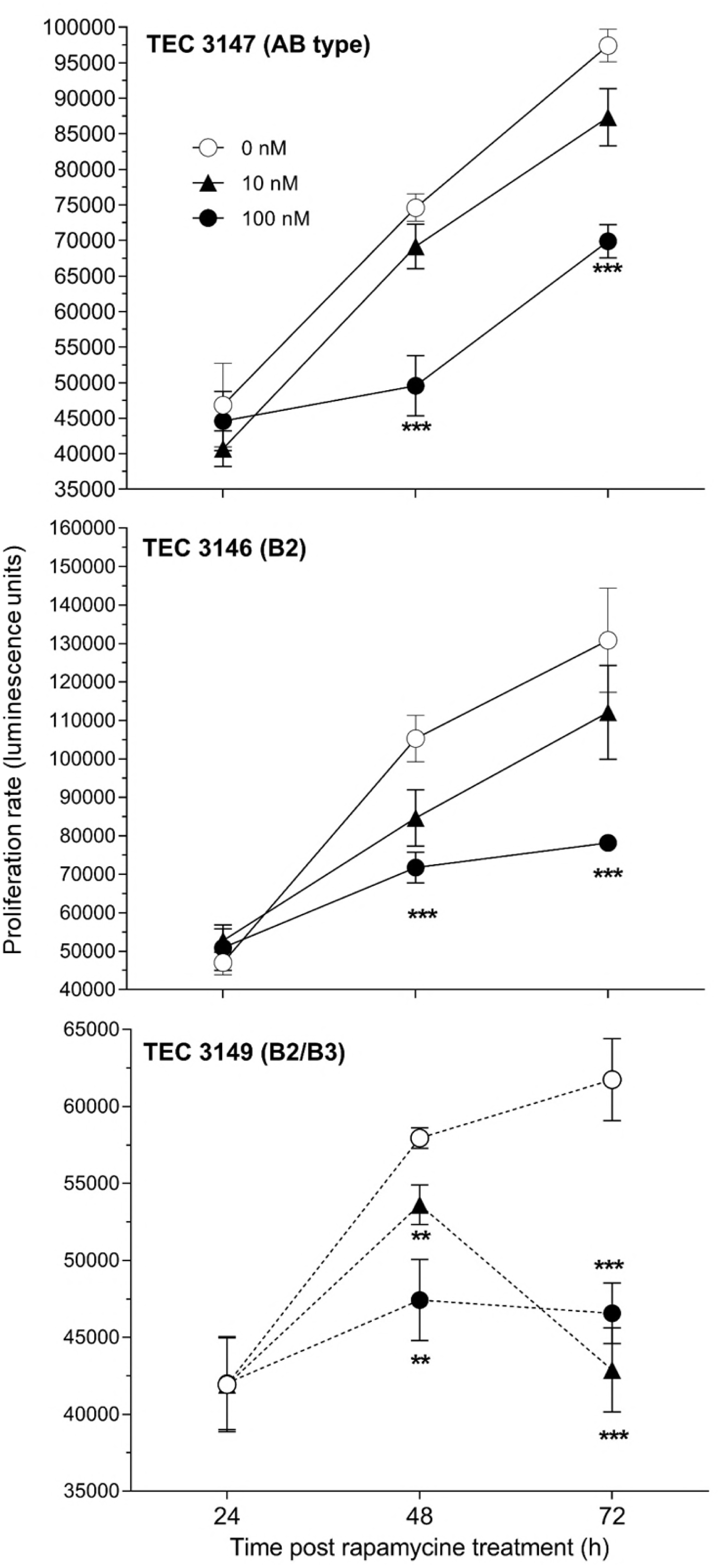
Inhibition of thymoma-derived epithelial cells upon rapamycin treatment. Proliferation rate has been measured in epithelial cells derived from AB (#3147), B2 (#3146) and B2/ B3 (#3149) thymomas after 24, 48 and 72 hours of culture with 0 nM (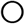), 10 nM (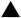) or 100 nM (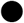) rapamycin. Statistical significance (α= 0.05) with t test; ** <0.02; *** < 0.001

We controlled the inhibition of mTOR and phospho-mTOR in cells treated for 24 hours with 100 nM rapamycin by measuring the level of protein expression and confirmed that they were significantly reduced after 48 hours with rapamycin (Fig 6).

**Fig 6.**
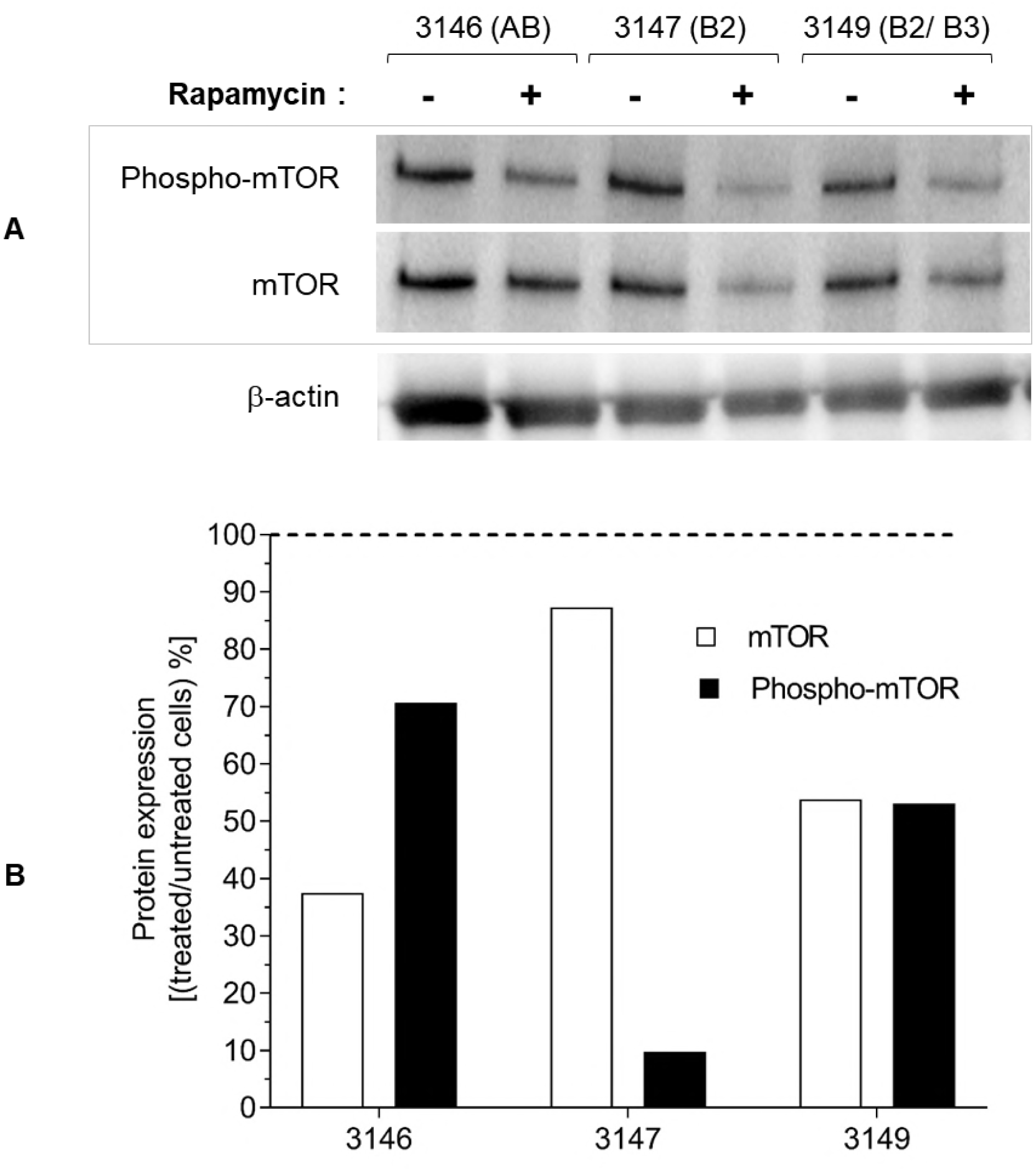
Inhibition of mTOR and phospho-mTOR expression upon rapamycin treatment of thymoma-derived cells. A. Expression of mTOR and phospho-mTOR proteins in thymoma-derived cells after 72h treatment with 100 nM rapamycin (+) or no rapamycin (−). B. Protein expression has been measured using and expressed as the ratio [treated/ untreated cells

## Discussion

Deciphering the molecular events in thymomas has remained a major challenge given the rarity of those tumors together with histological heterogeneity, precluding large genomic studies to be conducted, and the presence of lymphocytes intermixed with epithelial tumors cells in tissue specimens leading to potential misinterpretation of genomic features specifically associated with thymic carcinogenesis [27, 30, 31]. In this study, we report the feasibility of derivation of primary thymic epithelial cell cultures from type A, AB and B thymomas, their phenotypic and genetic characterization as well as the deregulation of the Akt-mTOR pathway and its impact on cell proliferation.

Previous reports of derivation of cells from thymic epithelial tumors have been made available. Most cell lines were obtained from thymic carcinoma specimens, with limited or more comprehensive subsequent molecular characterization. Besides PI3K regulatory subunits mutations [24], copy number gain of the anti-apoptotic molecule BCL2 was observed at comparative genomic hybridization of such cell lines, while *in vitro* siRNA knockdown reduced cell proliferation, and in vivo exposure to a pan-BCL2 inhibitor led to an inhibition of xenograft growth, *via* a mechanism involving the PI3K/AKT/mTOR pathway [23]. Exposure of thymic carcinoma cells to HSP90 inhibitors led to cell cycle arrest and apoptosis, and blocked invasiveness, through the downregulation of HSP90 oncogenic clients, including insulin-like growth factor 1 receptor (IGF-1R), a transmembrane tyrosine kinase receptor frequently overexpressed in thymic carcinomas, CDK4, and PI3K/Akt [32]. Taken together, these data were of significant therapeutic relevance: while pictilisib is mostly developed in breast cancers, which more frequently harbor PI3K alterations, phase II trials dedicated to thymic epithelial tumors were conducted with the IGF-1R inhibitor cixutumumab [33], the mTOR inhibitor everolimus [34] reporting on clinical antitumor activity in advanced, refractory cases. The IU-TAB-1-cell line was established from type AB thymoma, with phenotypic and molecular profiling but limited information of derivation protocol and success rate, and subsequent analysis of molecular pathways of interest [35].

In this study, we were able to derive primary thymic epithelial cells from all thirteen samples immediately after tumor removal, and successful expansion in culture. We provide the community with a reliable protocol that worked not only for thymic carcinomas but also for thymomas (data not shown), which are known to have lower proliferation index associated with slow growth and better outcome. We also show the feasibility of pathway analysis on those cells, including in vitro exposure to mTOR inhibitor, providing with a major tool to validate finding of high-throughput tissue analyses obtained on tissue specimens.

In our study, we found that the Akt-mTOR pathway was activated in thymic epithelial tumor cells derived from type A, AB, and B thymomas specimens, with correlation with corresponding tissue specimens; proliferation of those cells was significantly reduced after exposure to rapamycin through decrease of mTOR phosphorylation. We identified for the first time PI3KCA mutation in a type B2/B3 thymoma, which may be one mechanism of Akt-mTOR pathway, among others [36]. PI3K activation was also reported to be related to overexpression of a microRNA cluster on chr19q13.42 in type A and AB thymomas, observed in IU-TAB1 cell line [37]. This alteration was also observed in the cohort of The Cancer Genome Atlas [38]. Interestingly, only one of our cases harbored a GTF2I mutations, that was associated at RPPA analysis in this cohort, with lower expression of the apoptosis, cell cycle, DNA damage response, hormone receptor signaling, breast hormone signaling, RAS/MAPK, RTK, and TSC/mTOR pathways [38]. Our findings related to the activation of the Akt-mTOR pathway in a majority of specimens from patients with thymomas are of significant clinical relevance, given the recent results of a phase II study of everolimus in advanced thymic epithelial tumors, reporting on a disease control rate of 88%, with median progression-free survival of 10.1 months and median overall survival of 25.7 months [34]. Everolimus is currently available and may represent an off-label option for refractory tumors [3].

Ultimately, our data enhance the potential role of thymic epithelial cells derived from tissue specimens for an *in vitro* exploration of molecular abnormalities specific to thymic carcinogenesis. This may be relevant in a research setting to assess the value of molecular alterations observed at high-throughput genomic profiling, and develop in vivo models, but also to develop approaches for precision medicine strategies at the patient individual level.

## Bibliography

1. de Jong WK, Blaauwgeers JL, Schaapveld M, Timens W, Klinkenberg TJ, Groen HJ . Thymic epithelial tumours: a population-based study of the incidence, diagnostic procedures and therapy. Eur J Cancer. 2008 Jan;44(1):123–30. Pubmed PMID: 18068351.

2. Marx A, Strobel P, Badve SS, Chalabreysse L, Chan JK, Chen G, et al. ITMIG consensus statement on the use of the WHO histological classification of thymoma and thymic carcinoma: refined definitions, histological criteria, and reporting. J Thorac Oncol. 2014 May;9(5):596–611. Pubmed PMID: 24722150. Epub 2014/04/12.

3. Girard N, Ruffini E, Marx A, Faivre-Finn C, Peters S, Committee EG . Thymic epithelial tumours: ESMO Clinical Practice Guidelines for diagnosis, treatment and follow-up. Ann Oncol. 2015 Sep;26 Suppl 5:v40–55. Pubmed PMID: 26314779.

4. Masaoka A, Monden Y, Nakahara K, Tanioka T . Follow-up study of thymomas with special reference to their clinical stages. Cancer. 1981 Dec 01;48(11):2485–92. Pubmed PMID: 7296496.

5. Chen G, Marx A, Chen WH, Yong J, Puppe B, Stroebel P, et al. New WHO histologic classification predicts prognosis of thymic epithelial tumors: a clinicopathologic study of 200 thymoma cases from China. Cancer. 2002 Jul 15;95(2):420–9. Pubmed PMID: 12124843.

6. Regnard JF, Magdeleinat P, Dromer C, Dulmet E, de Montpreville V, Levi JF, et al. Prognostic factors and long-term results after thymoma resection: a series of 307 patients. J Thorac Cardiovasc Surg. 1996 Aug;112(2):376–84. Pubmed PMID: 8751506.

7. Safieddine N, Liu G, Cuningham K, Ming T, Hwang D, Brade A, et al. Prognostic factors for cure, recurrence and long-term survival after surgical resection of thymoma. J Thorac Oncol. 2014 Jul;9(7):1018–22. Pubmed PMID: 24926546. Epub 2014/06/14.

8. Berruti A, Borasio P, Gerbino A, Gorzegno G, Moschini T, Tampellini M, et al. Primary chemotherapy with adriamycin, cisplatin, vincristine and cyclophosphamide in locally advanced thymomas: a single institution experience. Br J Cancer. 1999 Nov;81(5):841–5. Pubmed PMID: 10555755. Pubmed Central PMCID: PMC2374302. Epub 1999/11/11.

9. Girard N, Lal R, Wakelee H, Riely GJ, Loehrer PJ. Chemotherapy definitions and policies for thymic malignancies. J Thorac Oncol. 2011 Jul;6(7 Suppl 3):S1749–55. Pubmed PMID: 21847058. Epub 2011/08/24.

10. Loehrer PJ Sr., Chen M, Kim K, Aisner SC, Einhorn LH, Livingston R, et al. Cisplatin, doxorubicin, and cyclophosphamide plus thoracic radiation therapy for limited-stage unresectable thymoma: an intergroup trial. J Clin Oncol. 1997 Sep;15(9):3093–9. Pubmed PMID: 9294472. Epub 1997/09/19.

11. Girard N, Merveilleux du Vignaux C . How large databases may impact clinical practices for rare tumors-postoperative chemotherapy in thymic malignancies. Journal of thoracic disease. 2016 Aug;8(8):1863–4. Pubmed PMID: 27619628. Pubmed Central PMCID: PMC4999727. Epub 2016/09/14.

12. Lemma GL, Lee JW, Aisner SC, Langer CJ, Tester WJ, Johnson DH, et al. Phase II study of carboplatin and paclitaxel in advanced thymoma and thymic carcinoma. J Clin Oncol. 2011 May 20;29(15):2060–5. Pubmed PMID: 21502559. Pubmed Central PMCID: PMC3107762. Epub 2011/04/20.

13. Loehrer PJ, Sr., Kim K, Aisner SC, Livingston R, Einhorn LH, Johnson D, et al. Cisplatin plus doxorubicin plus cyclophosphamide in metastatic or recurrent thymoma: final results of an intergroup trial. The Eastern Cooperative Oncology Group, Southwest Oncology Group, and Southeastern Cancer Study Group. J Clin Oncol. 1994 Jun;12(6):1164–8. Pubmed PMID: 8201378.

14. Loehrer PJ, Sr., Wang W, Johnson DH, Aisner SC, Ettinger DS, Eastern Cooperative Oncology Group Phase IIT. Octreotide alone or with prednisone in patients with advanced thymoma and thymic carcinoma: an Eastern Cooperative Oncology Group Phase II Trial. J Clin Oncol. 2004 Jan 15;22(2):293–9. Pubmed PMID: 14722038.

15. Palmieri G, Marino M, Buonerba C, Federico P, Conti S, Milella M, et al. Imatinib mesylate in thymic epithelial malignancies. Cancer Chemother Pharmacol. 2012 Feb;69(2):309–15. Pubmed PMID: 21710245. Epub 2011/06/29.

16. Wang Y, Thomas A, Lau C, Rajan A, Zhu Y, Killian JK, et al. Mutations of epigenetic regulatory genes are common in thymic carcinomas. Scientific reports. 2014 Dec 08;4:7336. Pubmed PMID: 25482724. Pubmed Central PMCID: 4258655.

17. Moreira AL, Won HH, McMillan R, Huang J, Riely GJ, Ladanyi M, et al. Massively parallel sequencing identifies recurrent mutations in TP53 in thymic carcinoma associated with poor prognosis. J Thorac Oncol. 2015 Feb;10(2):373–80. Pubmed PMID: 25299233.

18. Druey KM . Bridging with GAPs: receptor communication through RGS proteins. Science’s STKE : signal transduction knowledge environment. 2001 Oct 16;2001(104):re14. Pubmed PMID: 11604548.

19. Hirano A, Emi M, Tsuneizumi M, Utada Y, Yoshimoto M, Kasumi F, et al. Allelic losses of loci at 3p25.1, 8p22, 13q12, 17p13.3, and 22q13 correlate with postoperative recurrence in breast cancer. Clin Cancer Res. 2001 Apr;7(4):876–82. Pubmed PMID: 11309336.

20. Penzel R, Hoegel J, Schmitz W, Blaeker H, Morresi-Hauf A, Aulmann S, et al. Clusters of chromosomal imbalances in thymic epithelial tumours are associated with the WHO classification and the staging system according to Masaoka. Int J Cancer. 2003 Jul 01;105(4):494–8. Pubmed PMID: 12712440.

21. Starostik P, Greiner A, Schultz A, Zettl A, Peters K, Rosenwald A, et al. Genetic aberrations common in gastric high-grade large B-cell lymphoma. Blood. 2000 Feb 15;95(4):1180–7. Pubmed PMID: 10666188.

22. Khoury T, Arshad A, Bogner P, Ramnath N, Zhang S, Chandrasekhar R, et al. Apoptosis-related (survivin, Bcl-2), tumor suppressor gene (p53), proliferation (Ki-67), and non-receptor tyrosine kinase (Src) markers expression and correlation with clinicopathologic variables in 60 thymic neoplasms. Chest. 2009 Jul;136(1):220–8. Pubmed PMID: 19318677.

23. Petrini I, Meltzer PS, Zucali PA, Luo J, Lee C, Santoro A, et al. Copy number aberrations of BCL2 and CDKN2A/B identified by array-CGH in thymic epithelial tumors. Cell death & disease. 2012 Jul 19;3:e351. Pubmed PMID: 22825469. Pubmed Central PMCID: 3406591.

24. Alberobello AT, Wang Y, Beerkens FJ, Conforti F, McCutcheon JN, Rao G, et al. PI3K as a Potential Therapeutic Target in Thymic Epithelial Tumors. J Thorac Oncol. 2016 Aug;11(8):1345–56. Pubmed PMID: 27117832. Epub 2016/04/28. eng.

25. Kossai M, Duchemann B, Boutros C, Caramella C, Hollebecque A, Angevin E, et al. Antitumor activity in advanced cancer patients with thymic malignancies enrolled in early clinical drug development programs (Phase I trials) at Gustave Roussy. Lung Cancer. 2015 Sep;89(3):306–10. Pubmed PMID: 26160757. Epub 2015/07/15.

26. Zucali PA, De Pas T, Palmieri G, Favaretto A, Chella A, Tiseo M, et al. Phase II Study of Everolimus in Patients With Thymoma and Thymic Carcinoma Previously Treated With Cisplatin-Based Chemotherapy. J Clin Oncol. 2018 Feb 1;36(4):342–9. Pubmed PMID: 29240542. Epub 2017/12/15.

27. Wheler J, Hong D, Swisher SG, Falchook G, Tsimberidou AM, Helgason T, et al. Thymoma patients treated in a phase I clinic at MD Anderson Cancer Center: responses to mTOR inhibitors and molecular analyses. Oncotarget. 2013 Jun;4(6):890–8. Pubmed PMID: 23765114. Pubmed Central PMCID: 3757246.

28. Scorsetti M, Leo F, Trama A, D’Angelillo R, Serpico D, Macerelli M, et al. Thymoma and thymic carcinomas. Crit Rev Oncol Hematol. 2016 Mar;99:332–50. Pubmed PMID: 26818050.

29. Dagogo-Jack I, Shaw AT . Crizotinib resistance: implications for therapeutic strategies. Ann Oncol. 2016 Sep;27 Suppl 3:iii42–iii50. Pubmed PMID: 27573756. Pubmed Central PMCID: 5003168.

30. Girard N, Shen R, Guo T, Zakowski MF, Heguy A, Riely GJ, et al. Comprehensive genomic analysis reveals clinically relevant molecular distinctions between thymic carcinomas and thymomas. Clin Cancer Res. 2009 Nov 15;15(22):6790–9. Pubmed PMID: 19861435. Pubmed Central PMCID: 2783876.

31. Zucali PA, Di Tommaso L, Petrini I, Battista S, Lee HS, Merino M, et al. Reproducibility of the WHO classification of thymomas: practical implications. Lung Cancer. 2013 Mar;79(3):236–41. Pubmed PMID: 23279873. Pubmed Central PMCID: 3575111.

32. Breinig M, Mayer P, Harjung A, Goeppert B, Malz M, Penzel R, et al. Heat shock protein 90-sheltered overexpression of insulin-like growth factor 1 receptor contributes to malignancy of thymic epithelial tumors. Clin Cancer Res. 2011 Apr 15;17(8):2237–49. Pubmed PMID: 21372220.

33. Rajan A, Carter CA, Berman A, Cao L, Kelly RJ, Thomas A, et al. Cixutumumab for patients with recurrent or refractory advanced thymic epithelial tumours: a multicentre, open-label, phase 2 trial. The lancet oncology. 2014 Feb;15(2):191–200. Pubmed PMID: 24439931. Pubmed Central PMCID: PMC3954701. Epub 2014/01/21.

34. Zucali PA, De Pas T, Palmieri G, Favaretto A, Chella A, Tiseo M, et al. Phase II Study of Everolimus in Patients With Thymoma and Thymic Carcinoma Previously Treated With Cisplatin-Based Chemotherapy. J Clin Oncol. 2017 Dec 14:JCO2017744078. Pubmed PMID: 29240542. Epub 2017/12/15.

35. Gokmen-Polar Y, Sanders KL, Goswami CP, Cano OD, Zaheer NA, Jain RK, et al. Establishment and characterization of a novel cell line derived from human thymoma AB tumor. Lab Invest. 2012 Nov;92(11):1564–73. Pubmed PMID: 22926645.

36. Girard N. Chasing Therapeutic Targets in Thymic Malignancies: Finding Needles in the Haystack to Frame a Comprehensive Canvas? J Thorac Oncol. 2016 Aug;11(8):1197–200. Pubmed PMID: 27453162.

37. Radovich M, Solzak JP, Hancock BA, Conces ML, Atale R, Porter RF, et al. A large microRNA cluster on chromosome 19 is a transcriptional hallmark of WHO type A and AB thymomas. Br J Cancer. 2016 Feb 16;114(4):477–84. Pubmed PMID: 26766736. Pubmed Central PMCID: PMC4815766. Epub 2016/01/15.

38. Radovich M, Pickering CR, Felau I, Ha G, Zhang H, Jo H, et al. The Integrated Genomic Landscape of Thymic Epithelial Tumors. Cancer cell. 2018 Feb 12;33(2):244–58 e10. Pubmed PMID: 29438696. Epub 2018/02/14.

